# Absorption dipole effects on MINFLUX single molecule localization

**DOI:** 10.64898/2026.01.11.698872

**Authors:** Sjoerd Stallinga, Wenxiu Wang, Bernd Rieger

## Abstract

Single molecule fluorescence localization with minimum photon flux imaging (MINFLUX) can achieve localization precisions in the small nanometer range or better under suitable conditions. Potentially adverse conditions, such as a fixed fluorescence dipole or optical aberrations, that could cause systematic localization errors, have received little attention up to now. Here, we study these effects in simulation. We find that biases occur for fluorophores with a fixed absorption dipole tilted out of the imaging plane. These become larger (up to about 25% of the diameter of the circle spanned by the doughnut center positions) the larger the tilt angle gets. As a rule of thumb the spread in bias is smaller than 5 nm in case the dipole orientation is less than 30° out of plane for the typical case of a doughnut probing circle of diameter 100 nm. For freely rotating dipoles only the primary aberrations astigmatism and coma contribute to bias. This bias depends on the position of the fluorophore inside the circular probing area of MINFLUX and can be significantly larger than the localization precision. We show that increasing the number of measurements over the circle from a triangular to a hexagonal pattern is beneficial for reducing bias in all cases. Iterative shrinking of the probing area can eliminate the position dependent bias completely, but a strong dependence on dipole orientation of the bias at the center of the probing area remains.

## 1. Introduction

The introduction of MINFLUX has offered a breakthrough in localization precision for single molecule localization microscopy [1]. The technique is based on illumination with a doughnut shaped excitation beam. A sequence of illumination events, where the doughnut center is laterally shifted, gives rise to a fluorescence emission photon count that varies with the distance between the fluorophore and the doughnut center. A triangulation method is then applied to provide an estimate of the fluorophore’s position. The key strength of MINFLUX is that the precision scales with the size of the area over which the doughnut excitation beam is shifted, which can in principle be chosen arbitrarily small as long as the fluorophore remains inside the shift range. The MINFLUX methodology has been extended to 3D localization [2, 3], iteratively shrinking search area has been included [2, 4] for making more efficient use of the photon budget, and adapted for widefield detection by using spatially periodic illumination patterns [5–8]. The technique has also found its way to a wide range of applications such as tracking molecular motion on the sub-ms time scale [9], tracking motor protein dynamics [10, 11], deep tissue imaging [12], intramolecular distance measurements [13], structural changes in the Nuclear Pore Complex (NPC) [14] and RNA transport through the NPC [15, 16].

So far, the role of fundamental optics on image formation and statistical parameter estimation for MINFLUX has received little attention. He et al. [17] studied the impact of optical aberrations on MINFLUX localization, and reported on sensitivity for astigmatism and to a lesser extent coma. So far, no studies have been made on the impact of a possibly fixed absorption dipole orientation on MINFLUX. The dependence of conventional single molecule localization on the orientation of the emission dipole is well studied [18, 19], and imaging a fluorophore with a fixed but unknown emission dipole can lead to large biases in localization. The dependency on the emission dipole moment can be used to extend conventional localization microscopy to also image molecular orientation, using Point Spread Function (PSF) engineering [20, 21], polarized detection, excitation polarization modulation [22, 23], or a combination of these techniques (see refs. [24, 25] for recent reviews). As localization information in MINFLUX is extracted by scanning the excitation beam and point detection, any polarization or anisotropy effects of the emission are irrelevant, however, this is not true for the absorption process. The electric field distribution inside the doughnut spot 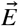interacts with the absorption dipole moment 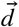to give an absorption PSF proportional to 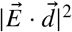, implying that deformations of the doughnut shape can be induced by a fixed absorption dipole moment of the fluorophore. Such deformations could have an impact on the localization of the fluorophore.

Here, we present a simulation study to address the impact of deformations of the doughnut excitation spot on MINFLUX localization, in particular the orientation of the fixed absorption dipole of the fluorophore under investigation. This study is similar in spirit to our earlier study of the impact of the orientation of a fixed emission dipole on localization in the context of conventional widefield imaging [18]. In section 2 we start from an analysis of the absorption PSF by vectorial optics, taking all effects of polarization and high NA into account. In section 3 we present a framework for Maximum Likelihood Estimation (MLE) for fitting the position of the absorber from data with Poisson noise. We apply the combination of physically exact PSF modeling and MLE fitting in a simulation setup for assessing precision (statistical spread) and accuracy (absence of bias) in section 4. Conclusions and outlook to follow-up work are presented in section 5.

## 2. Doughnut absorption PSF models

In Appendix A we derive expressions for the absorption PSF in case of a freely rotating absorption dipoles in the ideal aberration-free case:

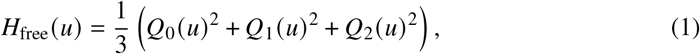

where *Q*_0_ (*u*), *Q*_1_ (*u*), and *Q*_2_ (*u*) are functions of the radial image plane coordinate *u* defined in the Appendix. Figure 1ab) show an example plot of the vector doughnut PSF for a freely rotating absorption dipole, computed for a wavelength *λ* = 520 nm, a Numerical Aperture NA = 1.45, where the doughnut is focused into a medium with refractive index *n* = 1.52. The free dipole absorption PSF is well approximated with a Gaussian doughnut PSF:

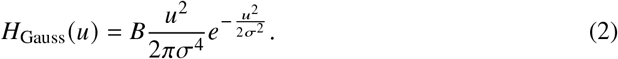

**Fig. 1.**
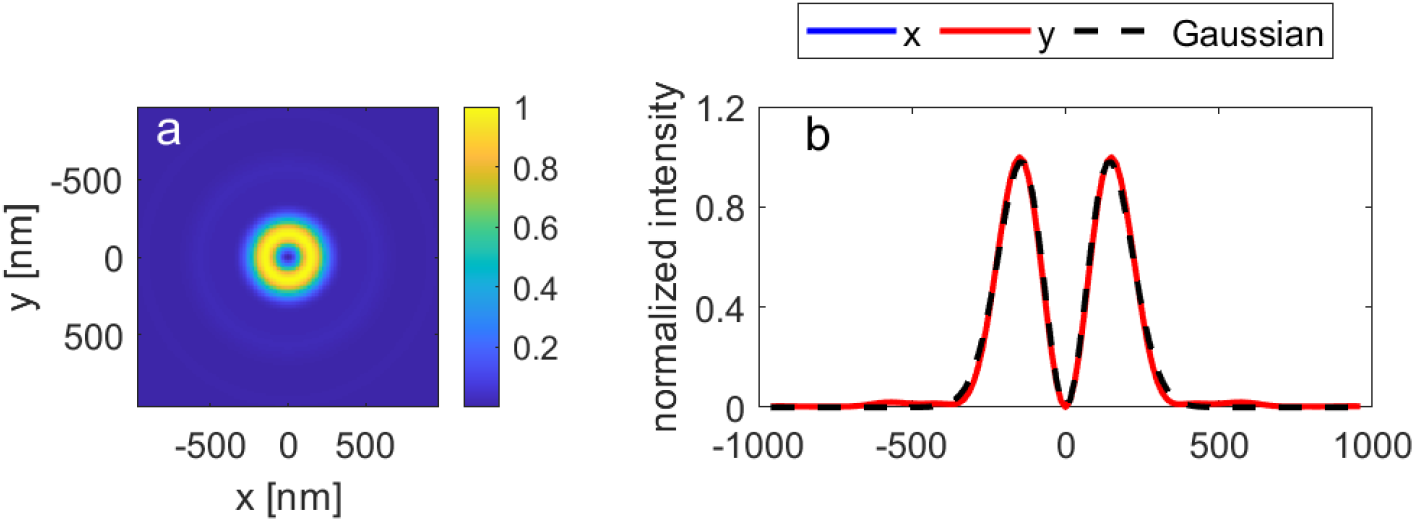
Vector doughnut PSF shape for freely rotating absorption dipoles. (a) Vector doughnut PSF for the nominal aberration-free case. (b) Cross-sections of vector doughnut PSF in comparison to Gaussian approximation.

For this example a width *σ* = 0.28*λ* NA and relative height *B* = 0.93 provides a good approximation to the doughnut PSF shape, where the largest mismatch is in the tail of the PSF outside the doughnut region (see Fig. 1b). We can use the approximate Gaussian doughnut PSF model as a simplified effective PSF model, similar in spirit to the use of the Gaussian PSF model in standard widefield imaging based emitter localization [18, 19].

For the case of fixed absorption dipoles and zero aberrations we also derive an expression for the absorption PSF in Appendix A:

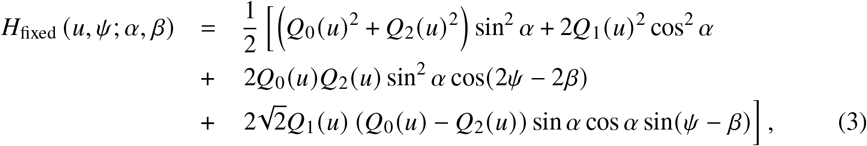

with *α* and *β* the polar and azimuthal absorption dipole angles, and where *ψ* is the azimuthal image plane coordinate. This expression for the PSF has different groups of terms with different dependency on the azimuthal angle *ψ*. The first two terms do not depend on *ψ*, but do show a crossover in doughnut shape from axial dipoles (*α* = 0) to lateral (in-plane) dipoles (*α* = *π* / 2). This crossover affects the type of intensity minimum at the optical axis. For axial dipoles the spot is fully determined by the function *Q*_1_ (*u*), which is proportional to *u*^2^ for small values of the (scaled) radial image plane coordinate. This implies that the doughnut minimum is no longer parabolic, but instead is 4th order. The third term makes the doughnut elliptical in the direction perpendicular to the projection of the dipole vector on the focal plane, an effect that scales from 0 for axial dipoles to a maximum for lateral dipoles because of the dependence on sin^2^ *α*. The fourth and final term is relevant for tilted dipoles, because of the scaling with sin *α* cos *α*, and makes the doughnut asymmetric, it pulls the center of gravity of the doughnut away from the focal point at *u* = 0 in the direction parallel to the projection of the dipole vector on the focal plane. The types of distortion of the doughnut shape with azimuthal dependence are qualitatively similar to the impact of astigmatism and coma [17], with the distinction that the intensity minimum is still zero in all cases, as opposed to the case of astigmatism. Figure 2 shows these deformations of the PSF shape. For comparison, the best fit Gaussian doughnut PSF for the freely rotating absorption dipole case is plotted along the cross-sections, showing how poor the approximation is in these cases.

**Fig. 2.**
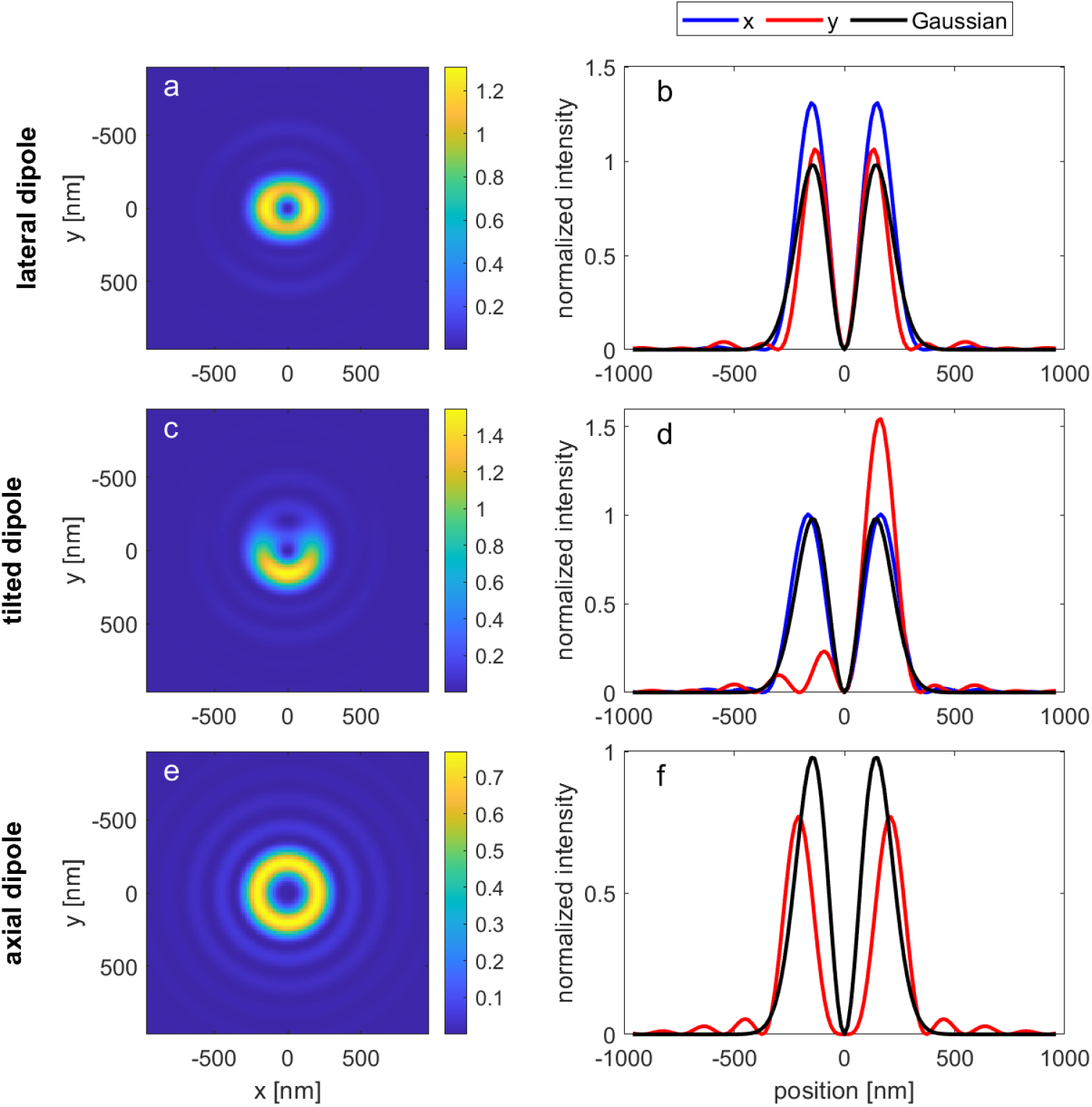
Vector doughnut PSF shape for fixed absorption dipoles. (a) Vector doughnut PSF shape for a lateral dipole (*α* = *π* / 2, *β* = 0). (b) Cross-sections of vector doughnut PSF for the lateral dipole case in comparison to Gaussian approximation. (cd) Vector doughnut PSF shape and cross-sections for a tilted dipole (*α* = *π* / 4, *β* = 0). (ef) Vector doughnut PSF shape and cross-sections for an axial dipole (*α* = 0).

## 3. Maximum Likelihood Estimation with effective Gaussian PSF model

An important design choice in implementing MINFLUX is the set of doughnut center coordinates that is used to probe the fluorophore position, the so-called Targeted Coordinate Pattern (TCP) [3]. We consider the implementation of MINFLUX where the fluorophore position is probed by *M*+1 positions: *M* on a circle of diameter *L* separated by 360 /*M* degrees each and one at the center of the circle. In particular we consider the triangular TCP case (*M* = 3), the original choice used in ref. [1] and applied as well in later studies [12, 26], and the hexagonal TCP case (*M* = 6), which is more often used in recent work on MINFLUX [11, 13]. We assume that the fluorophore is already inside the probing circle of diameter *L*. We label the *M* + 1 positions as:

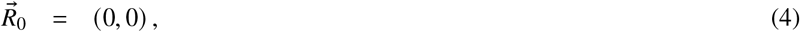

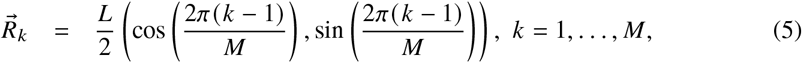

with the fluorophore position indicated as:

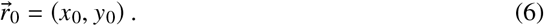

The expected photon counts for measurements *k* = 0, 1, 2, 3 are:

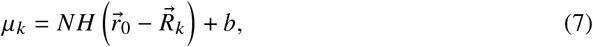

where 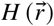 is the absorption PSF, *N* is the *total* signal photon count and *b* a constant background photon count per measurement. That means that the total number of background photons is (*M* +1)*b*.

The four unknown parameters *θ*_1_ = *x*_0_, *θ*_2_ = *y*_0_, *θ*_3_ = *N* and *θ*_4_ = *b* are estimated by optimizing the log-likelihood, given the Poisson statistics of the measurements as usual [1, 27]:

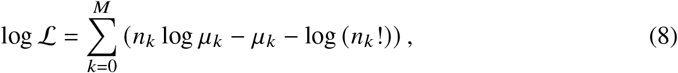

where *n*_*k*_ is the actually observed photon count for measurement *k*. In the numerical optimization it is helpful to use analytical derivatives of the log-likelihood w.r.t. the fit parameters:

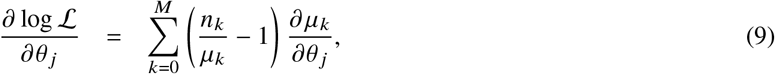

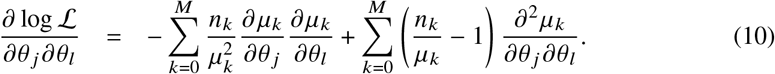

We have implemented the Levenberg-Marquardt method, where for the Hessian (matrix of second order derivatives), we use the approximation in which the actual photon numbers *n*_*k*_ are replaced by the expected photon numbers *μ*_*k*_. Of course, this does not alter the point of maximum likelihood, which is defined by the gradient being zero.

The expected photon counts *μ*_*k*_ and its derivatives depend explicitly on the assumed absorption PSF model. We use the approximate, and computationally efficient, Gaussian doughnut PSF model throughout the numerical simulation for fitting only:

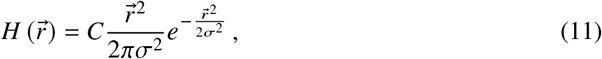

where the parameter *C* is a normalization factor to ensure that:

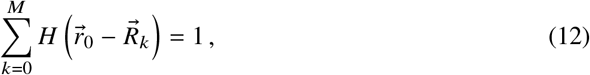

so that the parameter *N* corresponds to the expected total signal photon count. The Poisson rates can now be expressed as:

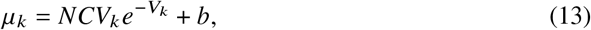

with:

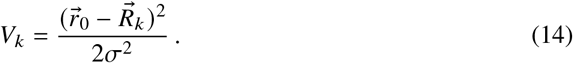

The normalization constant *C* is thus given by:

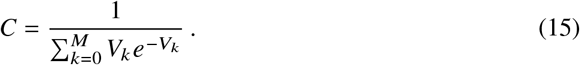

The first order derivatives of the Poisson rates *μ*_*k*_ follow as:

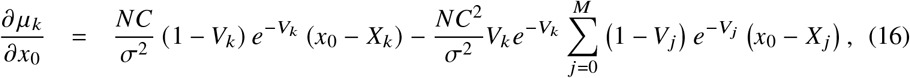

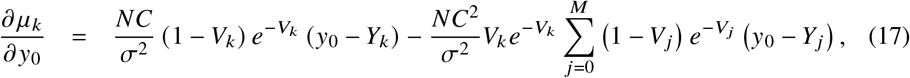

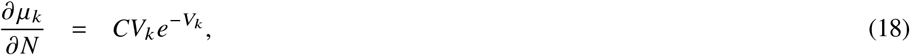

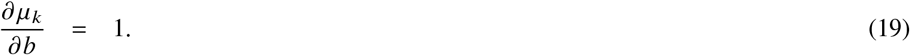

These expressions suffice for a numerical implementation of the Levenberg-Marquardt based MLE algorithm.

The Cramér Rao Lower Bound (CRLB) is found from the diagonal elements of the inverse of the Fisher matrix [27]. The Fisher matrix is evaluated according to:

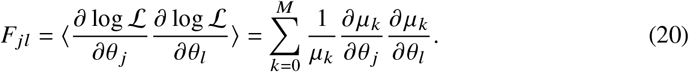

A concise analytical expression for the CRLB can be found for the case where the fluorophore is at the center doughnut position (*x*_0_ = *y*_0_ = 0), as derived in Appendix B, yielding the following estimation uncertainties:

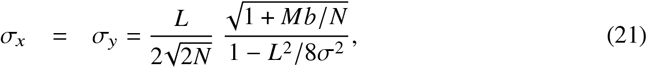

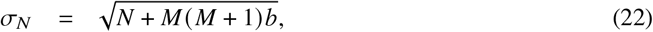

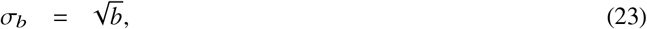

which differs from the expression for the localization uncertainty derived by Balzarotti et al. [1] in the dependence on background. This difference is due to a different parametrization of the background into a signal-to-background ratio in their case.

Initial values for the fluorophore coordinates *x*_0_ and *y*_0_ can be obtained from a phasor approach, similar to Martens et al. [28]. Using a parabolic approximation to the doughnut PSF we find that:

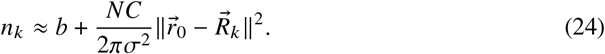

With this approximation, we compute the three linear weighted sums over the *M* measurement positions 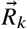 on the circle:

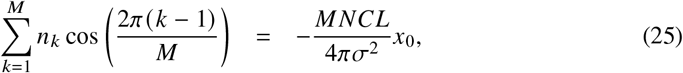

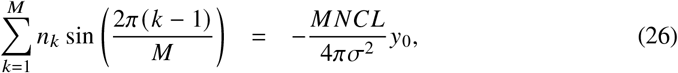

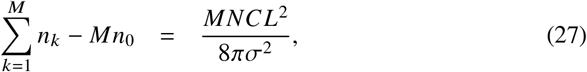

from which the coordinates follow by eliminating *MNC*/*πσ*^2^:

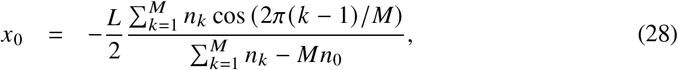

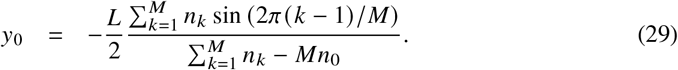

The key advantage of this phasor estimate is that the three weighted linear sums of photon counts are independent of the background photon count *b* and therefore the position estimate *x*_0_, *y*_0_ is not influenced by it. In addition, the recipe works for any value of the number *M* of doughnut positions on the circle of diameter *L*.

For the initial estimate of the signal photon count *N* and background photon count *b* we use a linear regression approach [29], i.e. given the above initial estimate of *x*_0_ and *y*_0_ we minimize:

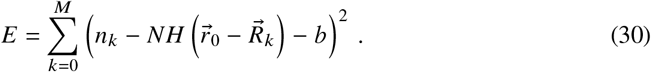

Values for *N* less than 1/10 times 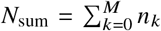 or larger than 10*N*_sum_ are clipped to this minimum or maximum value, respectively. Likewise, values for *b* less than −0.5 or larger than 0.5 times *N*_sum_ are clipped to this minimum or maximum value, respectively. With these initial value choices the numerical optimization procedure typically converges in 1 to 4 iterations, reaching on the order of 10^4^ fits/sec on CPU (on a consumer grade laptop).

## 4. Numerical assessment of accuracy and precision

We numerically evaluated the accuracy and precision by repeated localization of a known ground truth. Here the ground truth is simulated with the fully exact vectorial doughnut absorption PSF model, whereas the MLE is done with the simplified Gaussian doughnut PSF model. We take wavelength *λ* = 520 nm and NA = 1.45, where the doughnut is focused into a medium with refractive index *n* = 1.52. Typically, *S* = 200 optimizations of any set of ground truth parameter settings with different noise realizations are done. This choice for *S* is motivated by the practical trade-off between statistical precision (scales as 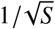) and computation time. The difference between the mean of the *S* outcomes and ground truth values of fit parameters measures the accuracy (bias), the standard deviation over the *S* outcomes measures the precision and is compared to the median CRLB of the *S* optimizations. We use the median CRLB as it is less sensitive to outlier fits. Please note that the CRLB captures only the statistical bound to an estimate, but no bias.

First, we generated different ground truth Poisson rates for the different doughnut positions (triangular TCP, *M* = 3) for a range of signal photon count values, background photon count values and positions within the MINFLUX search circle of diameter *L* for the case of freely rotating absorption dipoles in the aberration-free case. Figure 3 shows the mean of the fitted parameters, the standard deviation, and the CRLB across the MINFLUX search circle for *L* = 100 nm, total signal count *N* = 500 photons, and background *b* = 5 photons per measurement position. The localization bias vector (*B*_*x*_, *B*_*y*_), defined as the difference between the mean of the *S* = 200 localizations and the ground truth, shows that the estimation of the fluorophore position is virtually bias free (see Fig. 3a), in agreement with the close correspondence between the ground truth vector PSF model for freely rotating fluorophores and the Gaussian doughnut profile. The found bias values are very small and have a scattered profile across the FOV, pointing to incomplete averaging out of statistical errors in the sanple of size *S* = 200. The precision 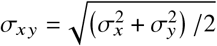 and 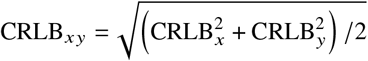 (Fig. 3c) match very well and indicate a very low (good) precision across the MINFLUX search circle, slightly increasing towards the edge defined by the circle through the three outer doughnut positions. The fitted values for the signal photon count *N* and for the background *b*, shown in Figs. 3d) and g) show a good match with the ground truth values in the center of the MINFLUX search circle and close to the positions of the three outer doughnut positions, but a small bias growing for fluorophore positions in between the three outer doughnut positions. The precision and CRLB for both *N* and *b* (Figs. 3e,f, h and i) correspond well and increase towards the regions in the MINFLUX search circle where a bias occurs. Figure 4 shows the corresponding results for a hexagonal TCP (*M* = 6). The fit performance is more uniform across the MINFLUX search circle, in particular for the fit of signal photon count *N* and background photon count per measurement *b*.

**Fig. 3.**
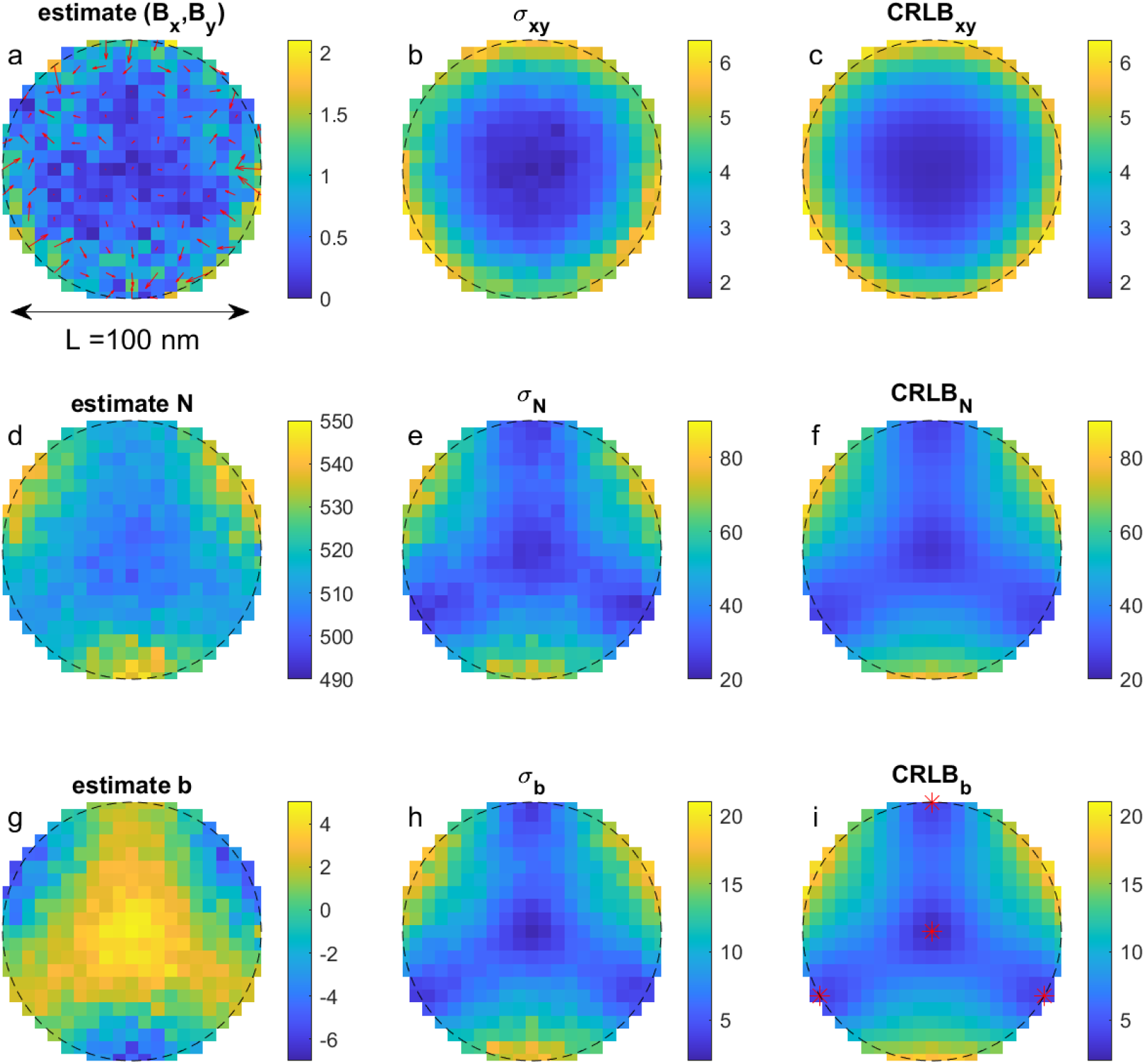
Gaussian doughnut fitting for freely rotating absorption dipole and nominal aberration-free case for triangular TCP (*M* = 3). (a) Bias vector (*Bx, By*). The color density plot indicates the magnitude, the arrow plot the orientation (the arrow length is 15× the actual bias vector to enhance visibility). (b) Localization precision averaged over *x* and *y* directions. (c) Median CRLB averaged over *x* and *y* directions. (d) Mean fitted signal photon count *N* (ground truth *N* = 500). (e) Fit precision of signal photon count. (f) Median CRLB of signal photon count. (g) Mean estimated background photon count *b* (ground truth *b* = 5). (h) Fit precision of background photon count. (i) Median CRLB of background photon count. The red asterisks indicate the positions of the four doughnut centers.

**Fig. 4.**
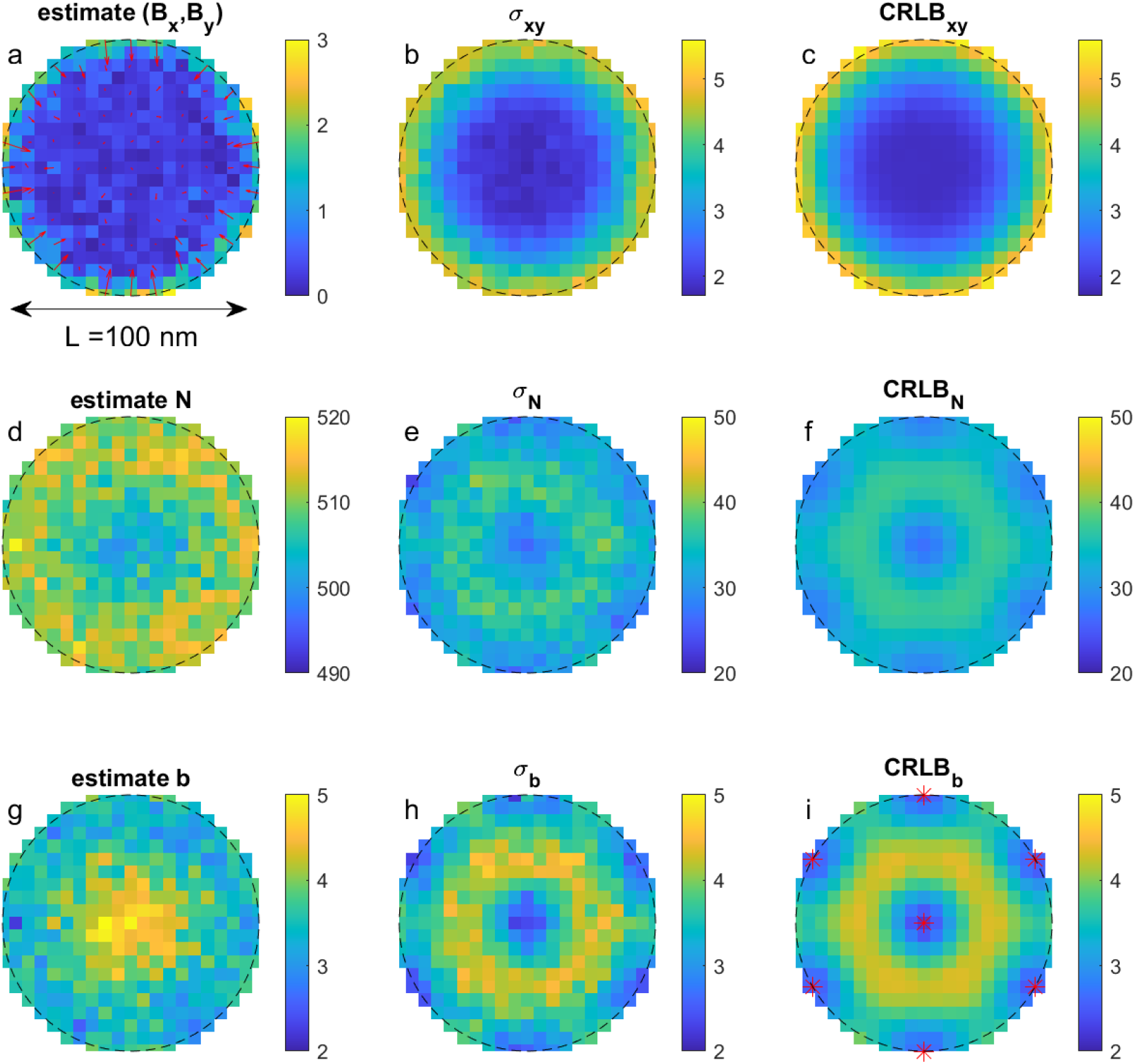
Gaussian doughnut fitting for freely rotating absorption dipole and nominal aberration-free case for hexagonal TCP (*M* = 6). (a) Bias vector (*Bx, By*) . The color density plot indicates the magnitude, the arrow plot the orientation (the arrow length is 15× the actual bias vector to enhance visibility). (b) Localization precision averaged over *x* and *y* directions. (c) Median CRLB averaged over *x* and *y* directions. (d) Mean fitted signal photon count *N* (ground truth *N* = 500). (e) Fit precision of signal photon count. (f) Median CRLB of signal photon count. (g) Mean estimated background photon count *b* (ground truth *b* = 5). (h) Fit precision of background photon count. (i) Median CRLB of background photon count. The red asterisks indicate the positions of the seven doughnut centers.

We further investigated the impact of position in MINFLUX search circle, size of the MINFLUX search circle *L*, signal photon count *N* and background photon count *b* on the localization bias and precision. We computed the bias vector (*B*_*x*_, *B*_*y*_), localization precision values *σ*_*x*_ and *σ*_*y*_, and median CRLB values CRLB_*x*_ and CRLB_*y*_ over the *S* = 200 independent noise realizations for a given ground truth setting of fluorophore coordinates, parameter *L* and signal and background photon count *N* and *b*. Next, we average the outcomes over a disk of radius *R* in the MINFLUX search circle, giving the average and the spread of the bias:

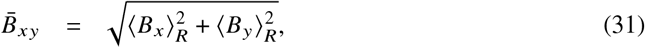

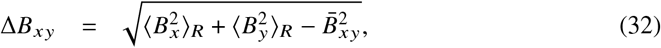

where *⟨*…*⟩*_*R*_ indicates the average over ground truth fluorophore positions within a disk of radius *R* from the center measurement. The average precision 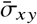 and average 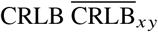 are defined similar to 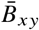. Figure 5a shows these quantities as a function of the radius *R* of the disk inside the MINFLUX search circle for the triangular TCP case (*M* = 3). The average precision grows somewhat from a value just below 2 nm to a value just above 3 nm as *R* increases from zero to the maximum value *L* / 2 = 50 nm. Figure 5b shows the impact of signal photon count *N* and background *b* on precision and CRLB. We observe the expected decrease with *N*, closely following the -1/2 slope, with background deteriorating the precision at lower signal photon counts. The precision reaches the CRLB for all but the lowest signal photon count values. Figure 5c shows the impact of the size parameter *L*, confirming the scaling of precision with *L*, the key strength of MINFLUX. The similar simulation for the hexagonal TCP case gave substantially the same outcome, only now the total number of background photons is now somewhat larger, giving a bit worse performance for given values of *N* and *b*. Next, we investigated the impact of a fixed absorption dipole orientation on the fit outcome. We computed ground truth photon rates based on the fixed dipole absorption PSF model for different values of the polar and azimuthal dipole angles *α* and *β*. We took the same default parameters as before (*L* = 100 nm, *N* = 500, *b* = 5, *S* = 200). The outcome of the numerical simulation for the triangular TCP case is shown in Figure 6. For lateral dipoles (*α* = *π* / 2, Fig. 6a, b, and c) the impact on localization bias is virtually absent. There is an asymmetry in the absorption PSF shape, but twofold rotational symmetry is maintained, and the intensity minimum is still strictly zero. For tilted dipoles (*α* = *π*/4, Fig. 6d, e, and f) there is a mean bias of several nm, and the bias depends considerably

**Fig. 5.**
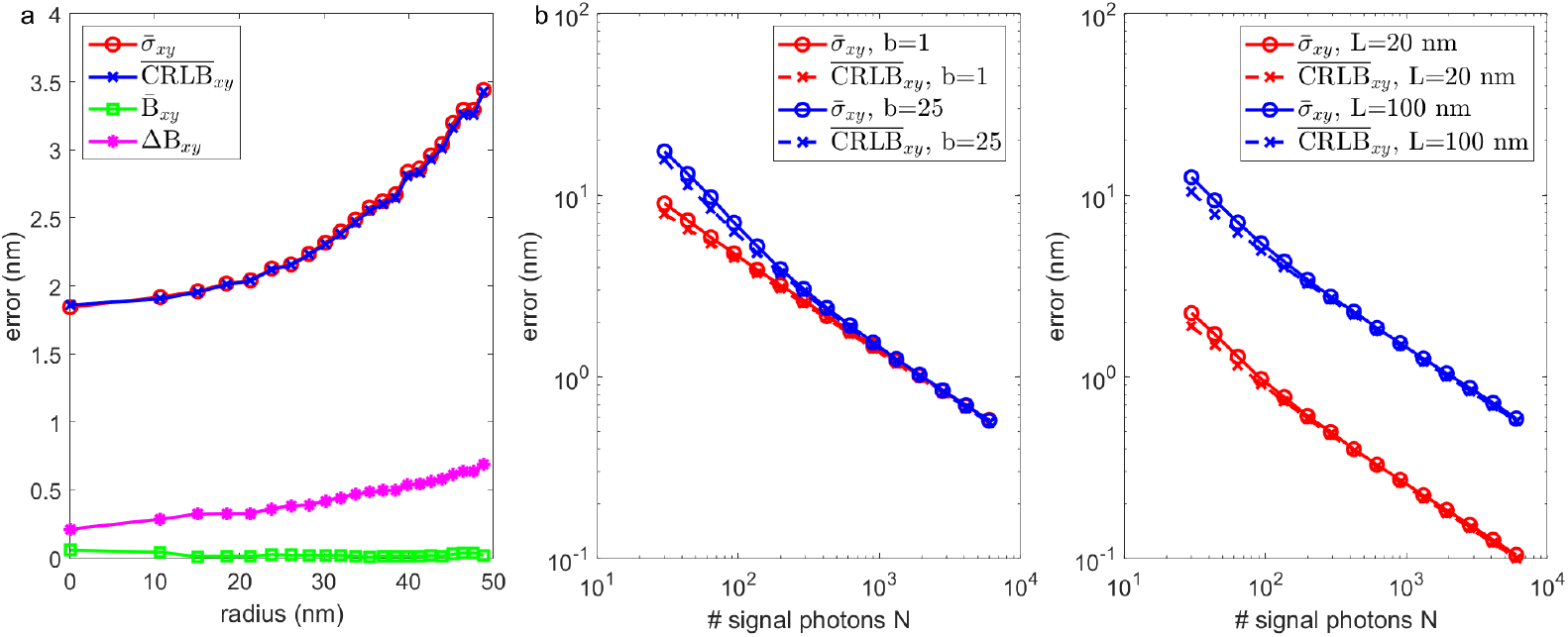
Impact of ground truth parameters on Gaussian doughnut fitting for freely rotating absorption dipole and nominal aberration-free case for triangular TCP (*M* = 3). (a) Average precision and CRLB, and average and spread of bias inside a disk of radius *R* in the MINFLUX search circle as a function of radius *R*. (b) Average precision and CRLB over a disk of radius *R* = 25 nm in the MINFLUX search circle as a function of signal photon count *N* for different values of the background photon count *b* (*L* = 100 nm). (c) Average precision and CRLB inside a disk of radius *R* = 25 nm in the MINFLUX search circle as a function of signal photon count *N* for different values of MINFLUX search circle diameter *L* (*b* = 5).

**Fig. 6.**
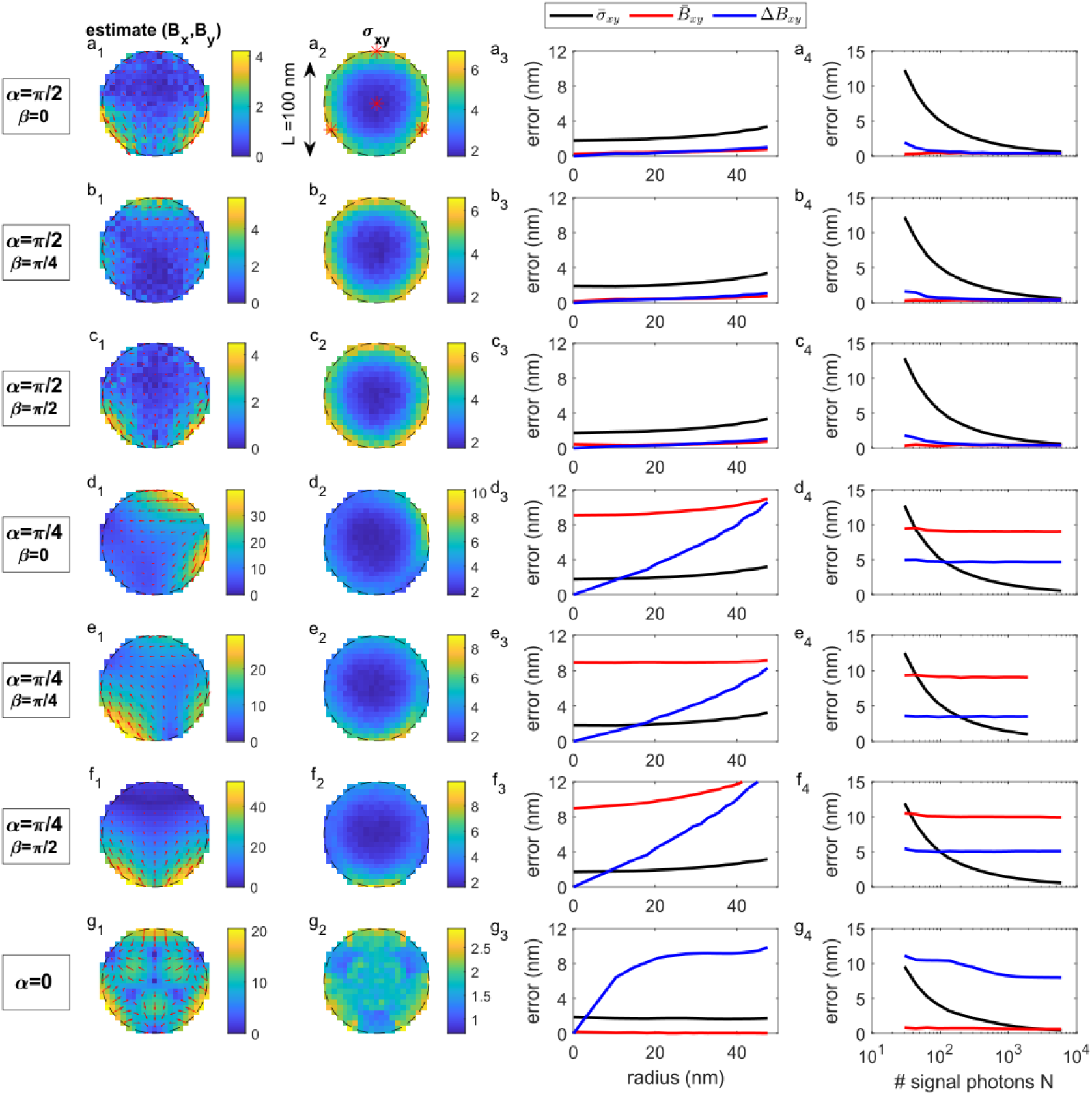
Gaussian doughnut fitting for absorption dipole with fixed orientation for triangular TCP (*M* = 3). Column 1 shows the bias vector across the MINFLUX search circle. The color density plot indicates the magnitude (in nm), the arrow plot the direction (the plotted length of the arrows is 15× the actually found bias vector for visibility purposes). Column 2 shows the localization precision across the MINFLUX search circle (color density plot in nm). Column 3 shows the average precision, average bias and spread of bias in a disk of radius *R* in the MINFLUX search circle as a function of radius *R* (*L* = 100 nm, *N* = 500, *b* = 5). Column 4 shows the average precision, average bias, and spread of bias in a disk of radius *R* = 25 nm as a function of photon count *N* (*L* = 100 nm, *b* = 5). (a) Results for lateral dipole *α* = *π*/2, *β* = 0 (polar angle *α*, azimuth angle *β*). (b) Results for lateral dipole *α* = *π*/2, *β* = *π*/4. (c) Results for lateral dipole *α* = *π*/2, *β* = *π*/2. (d) Results for tilted dipole *α* = *π*/4, *β* = 0. (e) Results for tilted dipole *α* = *π*/4, *β* = *π*/4. (f) Results for tilted dipole *α* = *π*/4, *β* = *π*/2. (g) Results for axial dipole *α* = 0.

on the fluorophore position, leading to a spread in bias that exceeds the precision outside a region of about 10 nm from the center doughnut position. Even though the PSF shape deformation is qualitatively similar to the shape deformation due to coma, the degree of deformation is much larger, leading to a stronger impact on localization inaccuracy. For axial dipoles (*α* = 0, Fig. 6g) we find the most intricate pattern of induced localization bias. There are 7 points with zero bias in the MINFLUX search circle (corresponding to the four doughnut center positions, and three points in between), with varying bias vectors in between. This results in an average bias over disks of radius *R* in the MINFLUX search circle that is near zero, as bias vectors have different orientations and cancel out, but with a spread that increases rapidly with radius *R*. The root problem of this erratic behaviour is the fourth-order intensity minimum of the doughnut absorption PSF for axial dipoles. As with the case of aberrations, the impact of the fixed dipole orientation on the precision is rather small, with precisions at the few nm level across the entire MINFLUX search circle. We again observed that the agreement with the CRLB is likewise good (data not shown).

Figure 7 shows the outcome of the same simulation for the hexagonal TCP case. Comparison to the triangular TCP case shows that the peak bias levels are reduced by a factor of about two. Also, the variation in bias with averaging disk radius *R* for the tilted dipole cases is strongly reduced, so that the maximum radius for which the bias spread in less than the precision is increased with a factor of about two to around 20 nm. For the axial dipole case the bias pattern is now rotationally symmetric, but the large bias spread remains, as this is rooted in the fourth-order intensity minimum.

**Fig. 7.**
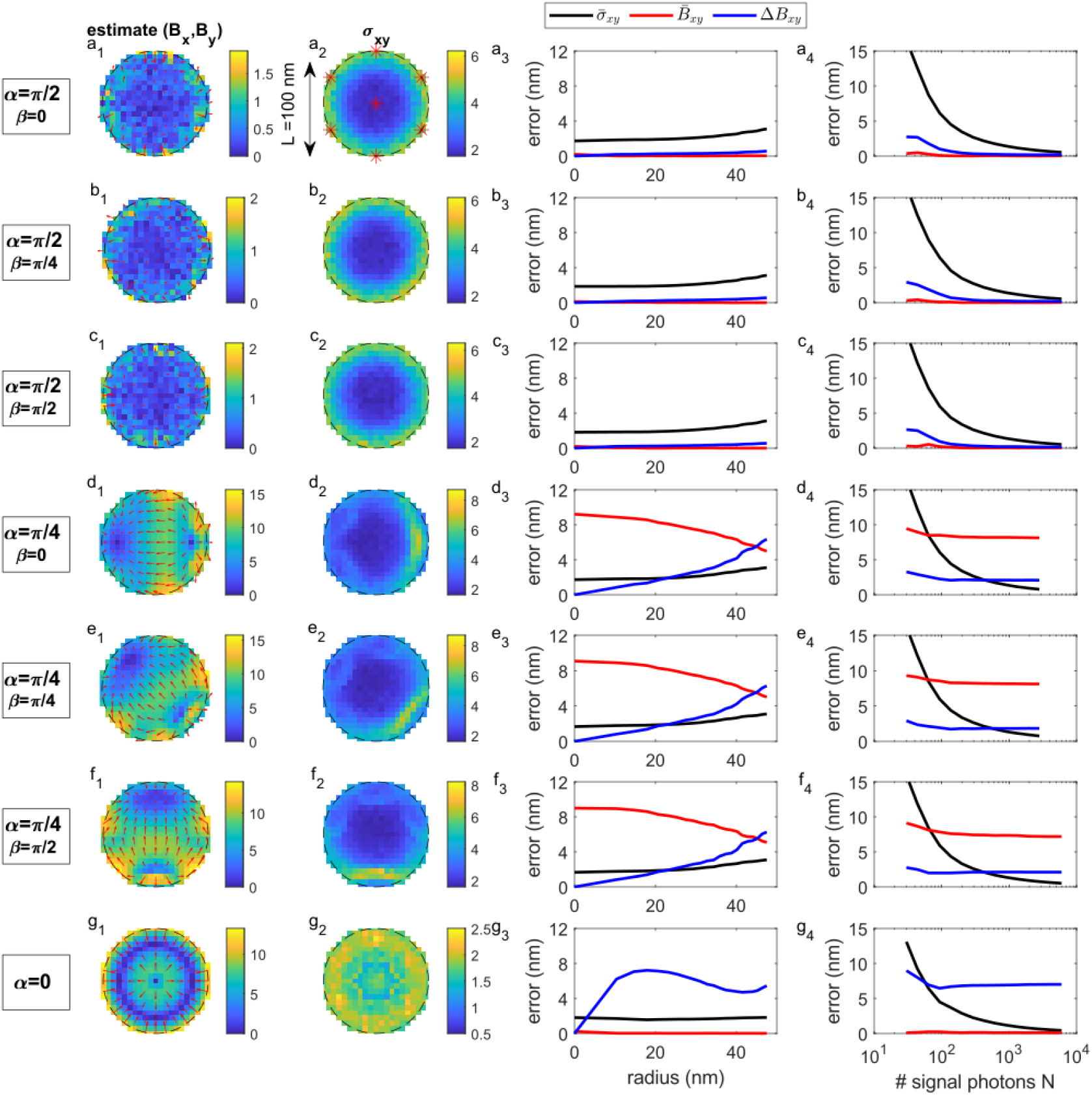
Gaussian doughnut fitting for absorption dipole with fixed orientation for hexagonal TCP (*M* = 6). Column 1 shows the bias vector across the MINFLUX search circle. The color density plot indicates the magnitude (in nm), the arrow plot the direction (the plotted length of the arrows is 15× the actually found bias vector for visibility purposes). Column 2 shows the localization precision across the MINFLUX search circle (color density plot in nm). Column 3 shows the average precision, average bias and spread of bias in a disk of radius *R* in the MINFLUX search circle as a function of radius *R* (*L* = 100 nm, *N* = 500, *b* = 5). Column 4 shows the average precision, average bias, and spread of bias in a disk of radius *R* = 25 nm as a function of photon count *N* (*L* = 100 nm, *b* = 5). (a) Results for lateral dipole *α* = *π*/2, *β* = 0 (polar angle *α*, azimuth angle *β*). (b) Results for lateral dipole *α* = *π*/2, *β* = *π*/4. (c) Results for lateral dipole *α* = *π*/2, *β* = *π*/2. (d) Results for tilted dipole *α* = *π*/4, *β* = 0. (e) Results for tilted dipole *α* = *π*/4, *β* = *π*/4. (f) Results for tilted dipole *α* = *π*/4, *β* = *π*/2. (g) Results for axial dipole *α* = 0.

The bias close to the center of the MINFLUX search circle is of importance for iterative MINFLUX where progressively smaller values of *L* are used. In case this is applied to fluorophores with a fixed yet unknown dipole orientation the bias at (*x*_0_, *y*_0_) = (0, 0) corresponds to the localization error. Figure 8a shows the fitted average localization error at (*x*_0_, *y*_0_) = (0, 0) as a function of the projected dipole orientation *d*_*x*_ = sin *α* cos *β* and *d*_*y*_ = sin *α* sin *β* for the triangular TCP case. For this center position in the MINFLUX search circle it appears that increasing *M* has no impact, for the hexagonal TCP case we find the same result. The bias peaks for a polar angle of around 10 degrees at a level as high as 25 nm for the parameter setting used (*L* = 100 nm, *N* = 500, *b* = 5). For larger polar angles the bias gradually decreases to a level around 1 nm for fully lateral absorption dipoles. For the range of polar angles between 60 and 90 degrees the variation in bias is around 5 nm. The bias dependence on dipole orientation may not be straightforward to detect. The largest bias occurs for dipoles with small polar angle, but these give rise to a doughnut shaped emission spot on the detector. In case of confocal detection with a narrow pinhole these spots may be suppressed or filtered out so that these near axial dipoles may be “not seen” in practice.

**Fig. 8.**
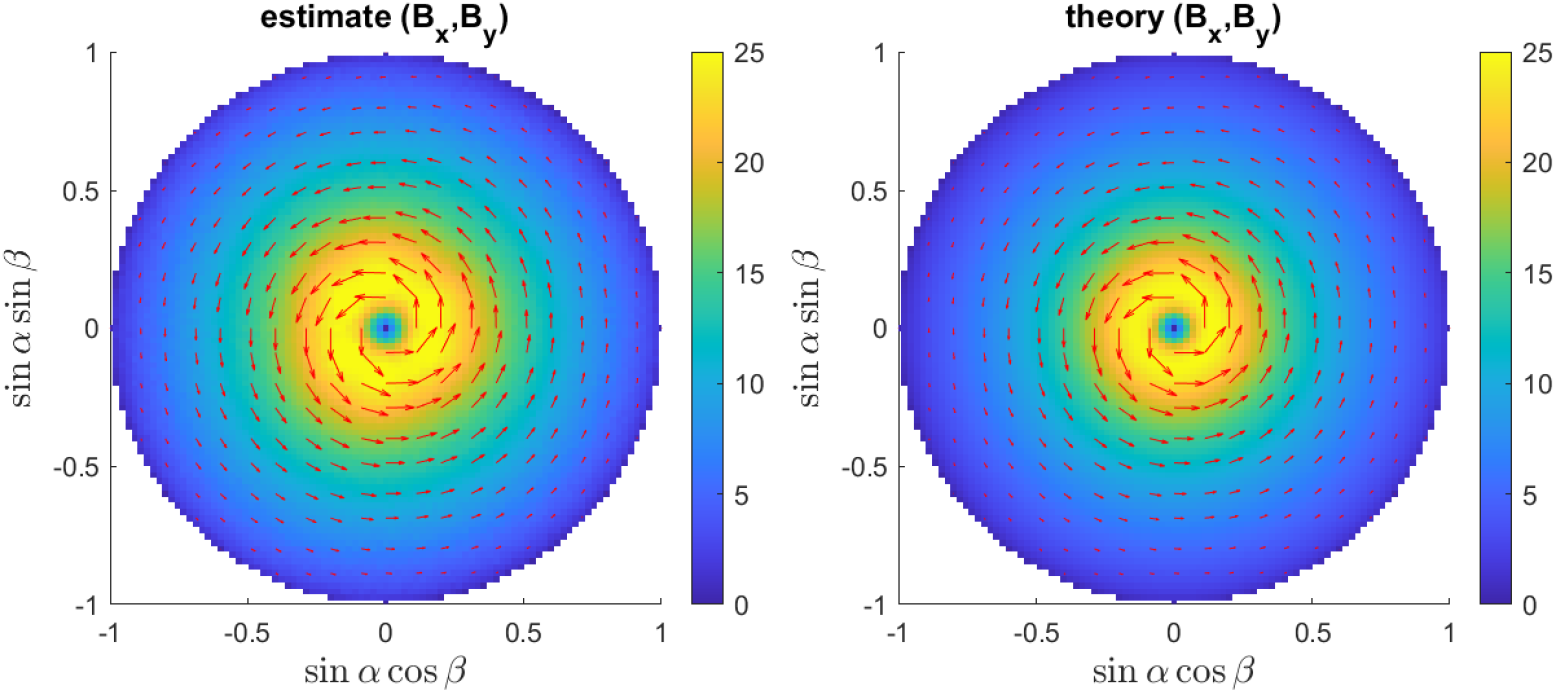
Dependence of Gaussian doughnut fitting bias on absorption dipole orientation. (a) Mean localization error (*Bx, By*) as a function of the projected dipole orientation *dx* = sin *α* cos *β* and *dy* = sin *α* sin *β*. The color density plot indicates the magnitude (in nm), the arrow plot indicates the orientation of the bias vector. (b) Phasor model prediction of the mean localization error (*Bx, By*) .

We compared the numerical results with a phasor model based estimate. For the ground truth case (*x*_0_, *y*_0_) = (0, 0) the analytical fixed dipole PSF model results in:

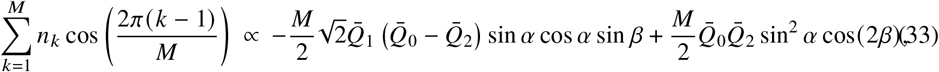

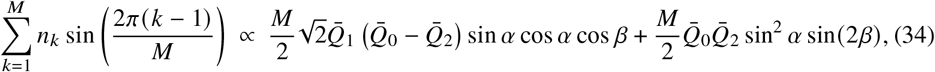

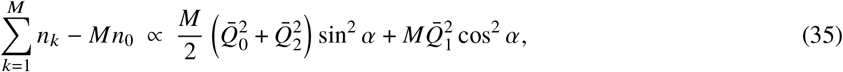

with 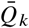 shorthands for *Q*_*k*_ (*L*NA/2*λ*) (*k* = 0, 1, 2). This gives rise to the apparent fluorophore coordinates:

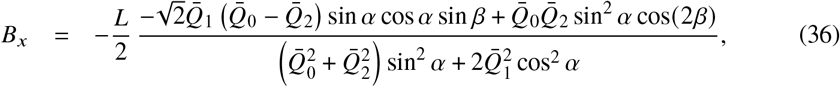

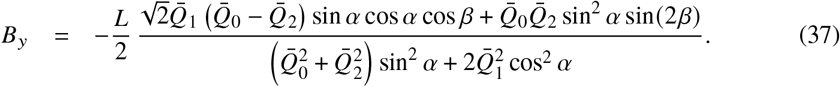

The numerical results match well with this model prediction (see Fig. 8b). The bias peak for fluorophores with relatively small polar angle *α* can be understood from the model if we consider that 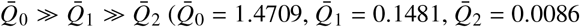 for the parameter setting used). Neglecting 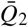 results in bias coordinates:

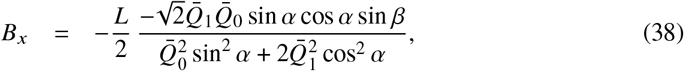

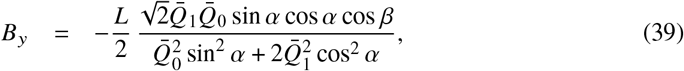

which for small *α* gives a bias magnitude that scales proportional to 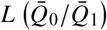 tan *α*. As 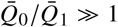 this bias magnitude grows linearly but steeply with *α*. The theoretical prediction is also independent of *M*, confirming the numerical result that the bias at (*x*_0_, *y*_0_) is independent of the choice of TCP (at least for the class of TCPs with doughnut centers organized on a circle). It turns out that the bias is proportional to the search circle diameter *L*, just like the precision. This implies that iteratively zooming in, reducing *L* to zero, will solve the bias problem in absolute terms but not in relative terms, as the ratio between bias and precision remains constant. This is somewhat similar to the case of conventional localization microscopy, where both precision and fixed dipole induced bias scale proportional to spot size *λ* / NA, implying for that case that a smaller spot size will reduce the bias problem in absolute terms, but not in relative terms.

Finally, we also revisited the study of He et al. [17] to compare the triangular and hexagonal TCP cases for the impact of the primary aberrations (astigmatism, coma, spherical aberration) on the performance of the fitter. We computed expected photon rates for the different doughnut positions for 5 different primary aberrations: two types of astigmatism (*A*_22_ and *A*_2−2_), two types of coma (*A*_31_ and *A*_3−1_), and spherical aberrations (*A*_40_). All aberrations are kept at the level of 36 m*λ* root-mean-square square (rms), i.e. at 50% of Maréchal’s tolerance criterion. We also modeled the fluorophore as a freely rotating absorption dipole. We computed the localization bias and precision across the MINFLUX search circle of diameter *L* = 100 nm for *N* = 500 and *b* = 5, based on a sample size *S* = 200. Figure 9 shows the results of the analysis for the triangular TCP case. For both types of astigmatism (Fig. 9a and b) we find a bias on the order of several nm, and an intricate dependence of bias on ground truth fluorophore position. As a consequence, the spread in the bias increase with patch radius *R*, and exceeds the precision for *R* larger than about 10 nm. We attribute this behaviour to the doughnut center which is not exactly zero for astigmatism (3% of peak intensity for this aberration setting). For both cases of coma (Fig. 9c and d) we also find a bias on the order of several nm, but now the bias is relatively constant across the MINFLUX search circle. This results in a spread in the bias that remains smaller than the precision up to large values of the patch radius *R*. For coma, the bias is induced by the asymmetry of the absorption PSF, while the intensity minimum remains zero. This results in the near-constant bias. For spherical aberration (Fig. 9e) the rotational symmetry of the absorption PSF is not broken, nor is there a non-zero intensity minimum. As a consequence the bias is virtually zero across the entire MINFLUX search circle. In all cases there is a surprisingly small impact on the localization precision, which hardly differs from the localization precision in the aberration-free case. We observed that the agreement with the CRLB is likewise good (data not shown). These findings are in agreement with the earlier study of He et al. [17]. Figure 10 shows the results of the analysis for the hexagonal TCP case. Compared to the triangular TCP case the bias patterns across the MINFLUX search circle for astigmatism and coma are much more symmetric, more uniform, and have a strongly reduced peak bias value (around threefold less for astigmatism, around twofold less for coma). This shows that MINFLUX with hexagonal TCP is much more robust against unknown aberrations than MINFLUX with triangular TCP. This agrees with the rationale for hexagonal TCP given in ref. [3].

**Fig. 9.**
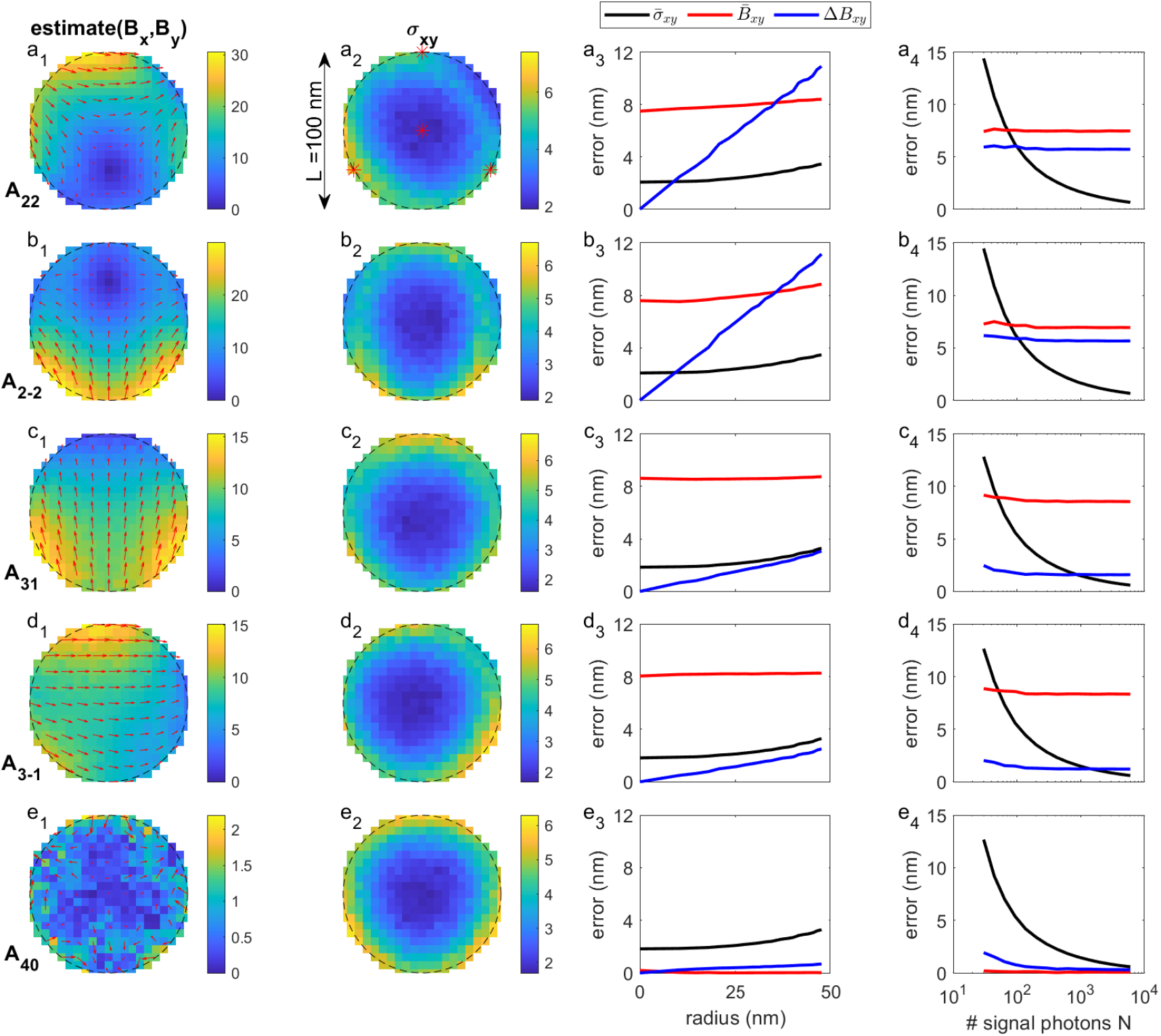
Gaussian doughnut fitting for freely rotating absorption dipole and different primary aberrations for triangular TCP (*M* = 3). Column 1 shows the bias vector across the FOV. The color density plot indicates the magnitude (in nm), the arrow plot the direction (the plotted length of the arrows is 15× the actually found bias vector for visibility purposes). Column 2 shows the localization precision across the MINFLUX search circle (color density plot in nm). Column 3 shows the average precision, average bias and spread of bias in a disk of radius *R* in the MINFLUX search circle as a function of radius *R* (*L* = 100 nm, *N* = 500, *b* = 5). Column 4 shows the average precision, average bias, and spread of bias in a disk of radius *R* = 25 nm as a function of photon count *N* (*L* = 100 nm, *b* = 5). (a) Results for horizontal/vertical astigmatism *A*_22_. (a)Results for diagonal astigmatism *A*_2−2_. (c) Results for horizontal coma *A*_31_. (d) Results for vertical coma *A*_3−1_. (e) Results for spherical aberration *A*_40_.

**Fig. 10.**
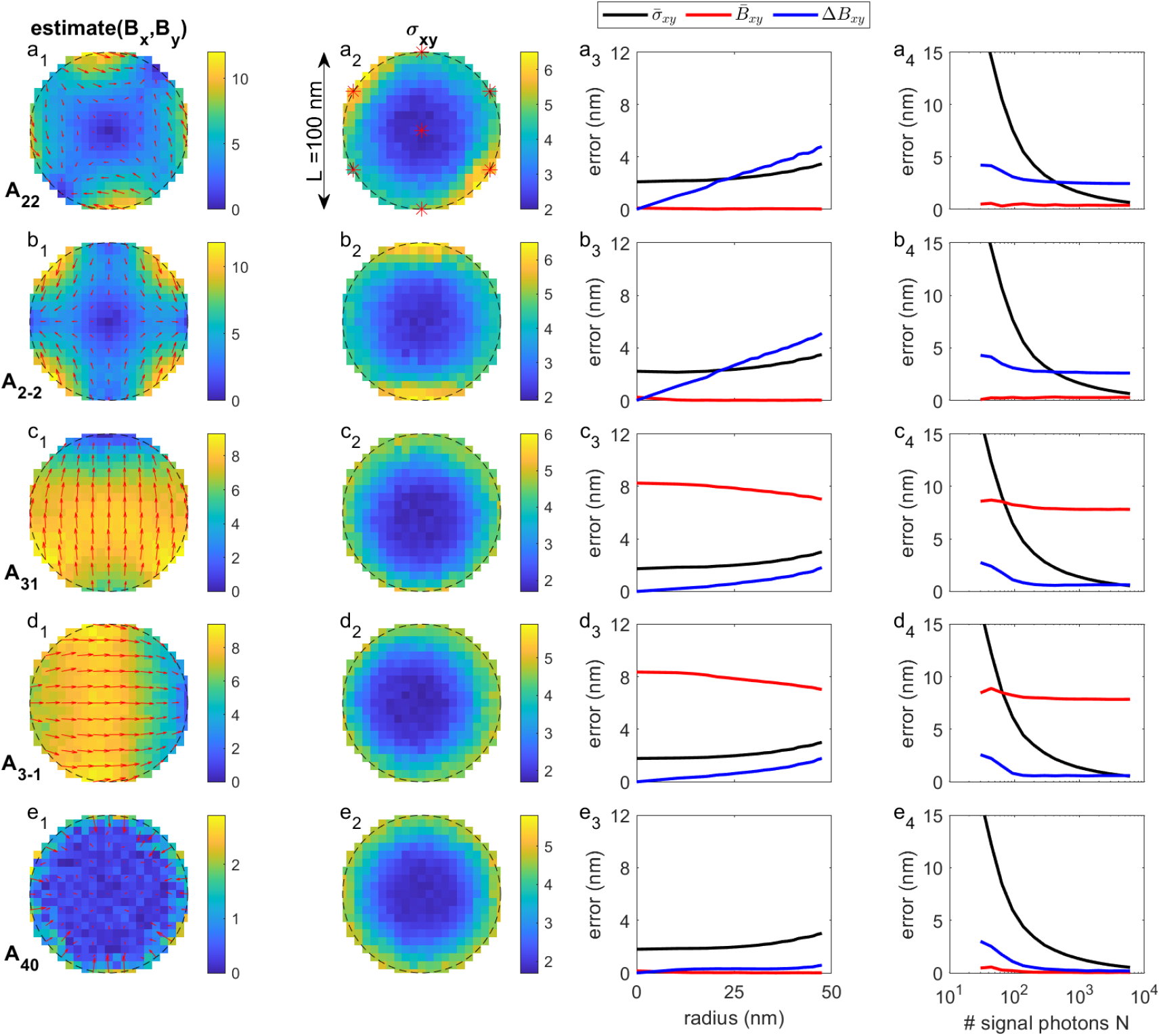
Gaussian doughnut fitting for freely rotating absorption dipole and different primary aberrations for hexagonal TCP (*M* = 6). Column 1 shows the bias vector across the FOV. The color density plot indicates the magnitude (in nm), the arrow plot the direction (the plotted length of the arrows is 15× the actually found bias vector for visibility purposes). Column 2 shows the localization precision across the MINFLUX search circle (color density plot in nm). Column 3 shows the average precision, average bias and spread of bias in a disk of radius *R* in the MINFLUX search circle as a function of radius *R* (*L* = 100 nm, *N* = 500, *b* = 5). Column 4 shows the average precision, average bias, and spread of bias in a disk of radius *R* = 25 nm as a function of photon count *N* (*L* = 100 nm, *b* = 5). (a) Results for horizontal/vertical astigmatism *A*_22_. (b) Results for diagonal astigmatism *A*_2−2_. (c) Results for horizontal coma *A*_31_. (d) Results for vertical coma *A*_3−1_. (e) Results for spherical aberration *A*,_40_.

## 5. Conclusion

In summary, we have studied the impact of possible mismatches between the ground truth vectorial absorption PSF model and MINFLUX localization using an effective Gaussian doughnut absorption PSF model. Overall, the localization precision closely follows the CRLB for a wide range of parameters (MINFLUX search circle diameter *L*, signal photon count *N*, background photon count *b*, ground truth fluorophore coordinates *x*_0_ and *y*_0_). We also proposed a phasor MINFLUX localization algorithm, which is computationally simple, insensitive to background, and which gives results close to the CRLB. In our MLE algorithm we use the phasor estimate for the initial value setting, which reduces the number of iterations to convergence to the range 1-4 depending on signal photon count.

We have focused in particular on the bias of MINFLUX localization in case of absorption PSF mismatches such as optical aberrations and a fixed absorption dipole orientation. We found that of the primary aberrations (astigmatism, coma, spherical aberration) astigmatism gives rise to a localization bias of several nm, that moreover is varying substantially with ground truth fluorophore position. For fluorophores with a fixed dipole orientation the ones with a small polar angle (close to axial orientation) result in a large bias that varies with ground truth fluorophore position. The root cause can be found in an imperfect doughnut minimum for the free and fixed dipoles, which is either non-zero (astigmatism) or of fourth-order (close to axial dipoles). Our study has been restricted to 2D MINFLUX. Analysis of the impact of optical aberrations and fixed absorption dipole effects in the context of 3D MINFLUX [2, 3, 30] requires a separate follow-up study.

It remains to be seen if the found localization bias effects are problematic in practice. For the case of aberrated absorption PSFs the iterative MINFLUX strategy [2, 4], with decreasing *L* values in each iterative step, the ground truth fluorophore position is increasingly confined to be close to the center doughnut position. In that case a constant bias is found, which can also be expected to be constant throughout the microscope’s Field Of View (if it is not too large the optical aberrations are typically constant). This gives rise to an overall shift in the entire image and is hence irrelevant. This line of reasoning is not applicable to the fixed absorption dipole case. Then the bias for fluorophores close to the center doughnut position varies with orientation over up to 25 nm, while different fluorophores that are localized with MINFLUX can have different (unknown) orientations. This could give rise to systematic errors in the distance between such dipoles in experiments such as by Sahl et al. [13]. These effects, however, are often unobserved in practice by the poor visibility of dipoles with a high tilt angle (nearly along the optical axis), because confocal detection of the doughnut shaped emission spots for such dipole orientations has a low efficiency. In addition, in aqueous solution fluorophores are free to rotate and different dipole orientations are measured even within one TCP scan, thus the here presented problem is strictly limited to (mostly) fixed fluorophores. Furthermore, it may be expected that in realistic circumstances fully fixed dipoles seldom occur, and that there is rather always some orientational flexibility left. The impact of partial orientational constraint could be taken into account by using a weighted sum of the free and fixed dipole absorption PSFs in the analysis [31].

We envision several methods that could reduce or eliminate the localization biases. Exploring these directions to eliminate biases will also offer the exciting opportunity to extend the MINFLUX methodology with estimation of molecular orientation. The first method is to simply image more than just the default four or seven doughnut center positions, in particular doughnut center positions at different *L* values. In this way the absorption PSF is sampled better and shape parameters could be extracted in addition to the regular four fit parameters (*x*_0_, *y*_0_, *N* and *b*). This could especially provide access to the possible non-zero intensity minimum and to the fourth-order component, which is informative on the polar dipole angle. In the framework of iterative MINFLUX [2] the data for such a method is already available, and a post-hoc overall estimation based on all individual iterative steps with shrinking *L* could eliminate the localization biases. Such a procedure could also be used as an alternative for pre-calibrating the absorption PSF [1]. The second method to address localization biases is to use a pixelated detector as in e.g. Image Scanning Microscopy (ISM) [32, 33] but now for measuring the shape of the emission PSF. Although this is not so informative on the fluorophore position as the MINFLUX recipe, it does provide information on possible optical aberrations and fixed dipole orientation. This approach requires that the absorption and emission dipoles are parallel, which is typically the case in practice. Exceptions do only occur when the excitation brings the fluorophore to an excited state other than the first single state (from where the fluorescence is emitted) [34]. The third method is to use polarized detection, with two detectors in each arm of a Polarizing Beam Splitter (PBS), and/or polarized excitation, by switching the polarization of the excitation laser. The fourth method is to engineer the absorption doughnut PSF, along the lines of [10, 35, 36], towards an absorption PSF shape with a more robust parabolic intensity minimum. In all cases more sophisticated modeling of the PSF, based on the physically motivated vectorial image formation as described in this paper by us, seems a necessity.

## Appendix A: Theory of vectorial doughnut PSF

The aim is to derive analytical theory for computing in-focus doughnut excitation PSF, following the well-known Richards-Wolf approach. It is convenient to work with scaled image and pupil coordinates:

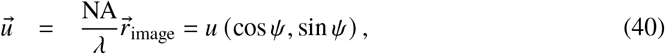

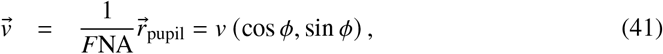

with *λ* the wavelength, NA the numerical aperture, and *F* the focal length of the objective lens. The advantage of using these diffraction units is that it maps the pupil onto the unit circle, and that all focal spot dimensions properly scale with *λ*/ NA. Each point in the pupil plane corresponds to an incoming plane wave towards focus with wavevector:

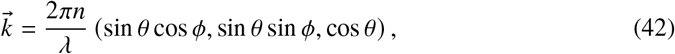

where *n* is the refractive index of the medium, and where the polar angle of incidence *θ* is related to the radial pupil coordinate *v* via Abbe’s sine condition:

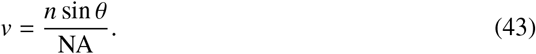

Figure 11 shows a layout of the focusing objective lens and the used coordinates in the pupil and focus plane.

**Fig. 11.**
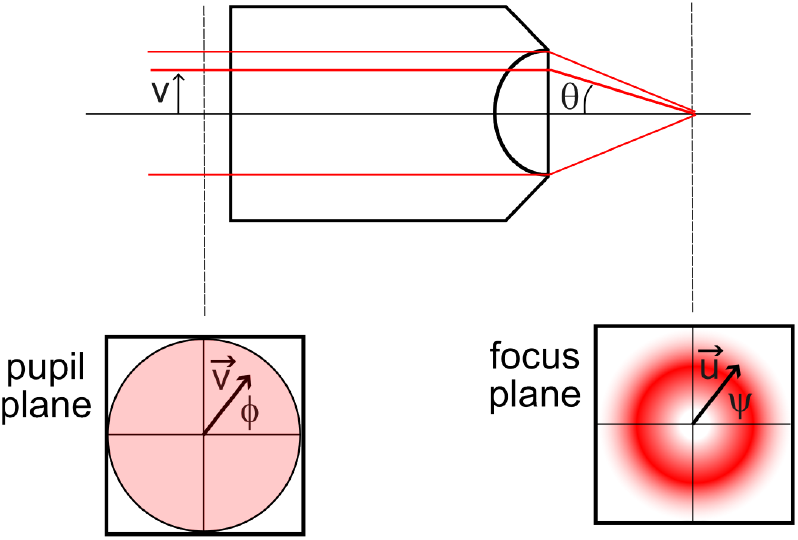
Schematic layout of focusing and coordinate conventions.

The components of the electric field in the image plane are given by the 2D Fourier Transformations:

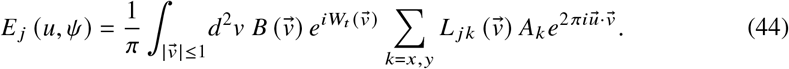

for *j* = *x, y, z*. Here, the *L* _*jk*_ 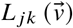 are the elements of the lens matrix:

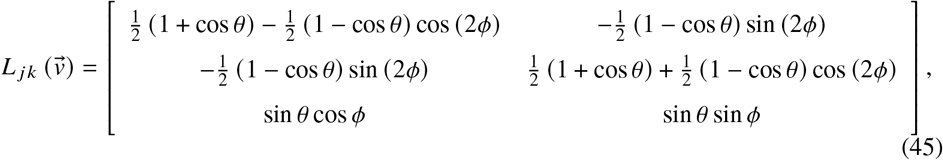

*B* 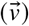 is the so-called aplanatic amplitude factor:

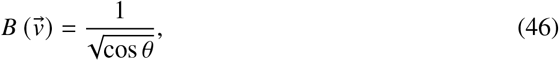

and W_*t*_ 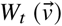 is the total pupil phase, which is equal to the sum:

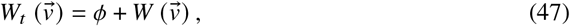

where the first term corresponds to the phase of the vortex phase plate and the second term represents the aberration function of the illumination optical system. Finally, the electric field components in the pupil plane are:

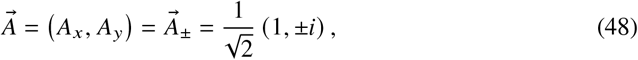

i.e. we consider the two circularly polarized polarization states as entrance polarization. Any entrance polarization state can be decomposed as the sum of these two states so this provides a complete description.

In case of aberrations, defocus, or imaging using evanescent as well as propagating waves, the diffraction integrals need to be evaluated numerically. For the ideal case without aberrations, for zero defocus, and for propagating wave imaging it turns out that analytical results are possible. In the following we will derive these results.

The product between the lens matrix and the entrance polarization vector results in:

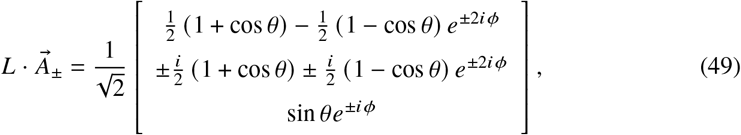

Next, we expand the Fourier exponential in a Bessel beam series:

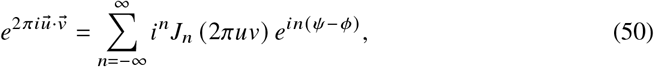

where the Bessel functions satisfy:

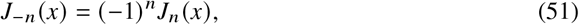

and evaluate the integral over the azimuthal pupil coordinate *ϕ* analytically using:

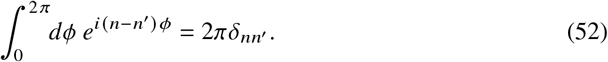

This results in an in-focus electric field with components:

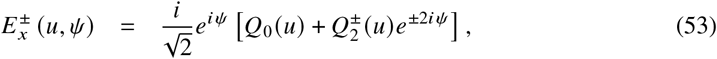

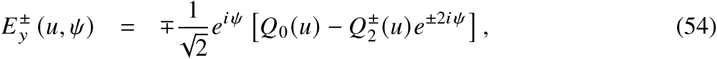

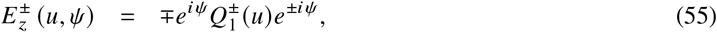

with:

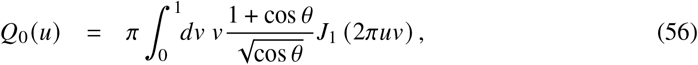

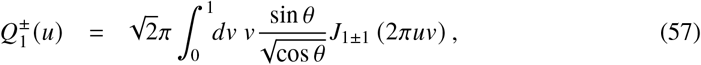

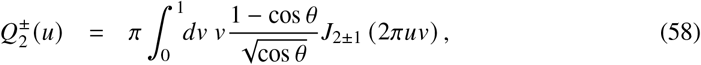

which are functions of the radial image plane coordinate *u*. Note that in evaluating these integrals the polar angle *θ* should be expressed in terms of the radial pupil coordinate *v* using the Abbe sine condition.

We can find more about the behaviour of the *Q* functions in different limiting cases. In the paraxial limit of low NA we may use that cos *θ ≈* 1 *− v*^2^NA2/*n*^2^ and we find:

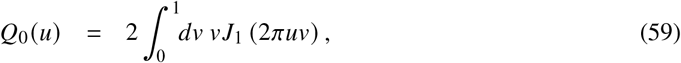

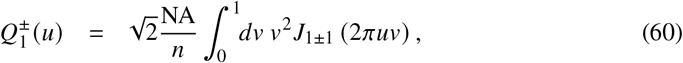

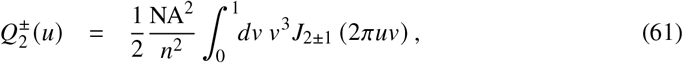

so that 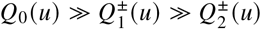, and we can ignore the contributions form 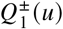 and 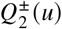. Close to the focal point *u ≪* 1 it holds that:

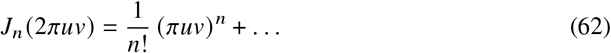

which implies that *Q*_0_(*u*) is then proportional to *u*, 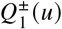 is proportional to *u*^2^ (for entrance polarization 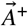) or a constant (for entrance polarization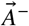), and 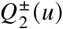 (*u*) is proportional to *u*^3^ (for entrance polarization 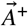) or to *u* (for entrance polarization 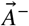). This implies that on the optical axis *u* = 0 all electric field components are zero for the left-handed circularly polarized entrance polarization 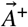. For the right-handed circularly polarized entrance polarization, however, there is a non-zero axial electric field component on the optical axis *u* = 0. It is therefore the first entrance polarization that gives rise to a doughnut spot with an exact zero at the focal point. In the following we will therefore only consider the 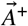case and use the abbreviations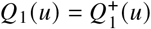, and 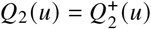 .

Defocus and spherical aberration have rotational symmetry in the pupil and add fringe structure to the doughnut spot outside the primary ring of the doughnut, but have limited impact on the qualitative shape of the doughnut minimum. This can be understood by a slight generalization of the above treatment for a rotationally symmetric aberration function *W v* with 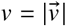. The field close to focus can now still be expressed in the functions *Q*_*n*_ *u* for *n* = 0, 1, 2, but now the integrals defining these functions include an additional phase factor exp (*i*W (*v*)) . In particular, it then still holds that *Q*_*n*_ (*u*)∼ *u*^*n*+1^ for small *u*. As a consequence, the doughnut minimum at *u* = 0 remains intact.

The in-focus absorption PSF for a fixed absorption can be expressed in terms of the functions *Q*_0_(*u*), *Q*_1_(*u*), and *Q*_2_(*u*) via:

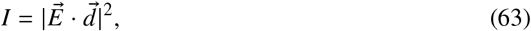

with the absorption dipole vector:

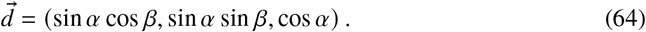

Here *α* is the polar dipole angle and *β* is the azimuthal dipole angle. In case of a freely rotating absorption dipole we need to average over all directions to get an absorption PSF:

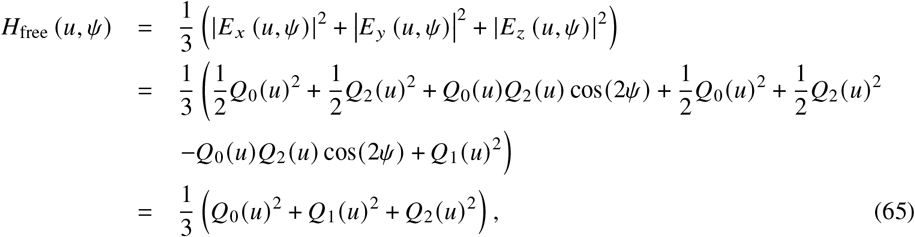

that turns out to be rotationally symmetric, i.e. independent of the azimuthal angle *ψ*. Next, we consider the fixed dipole case. Evaluating the inner product:

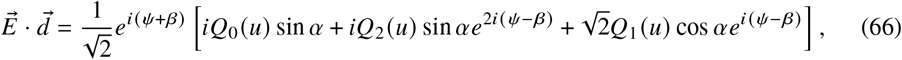

leads to the absorption PSF:

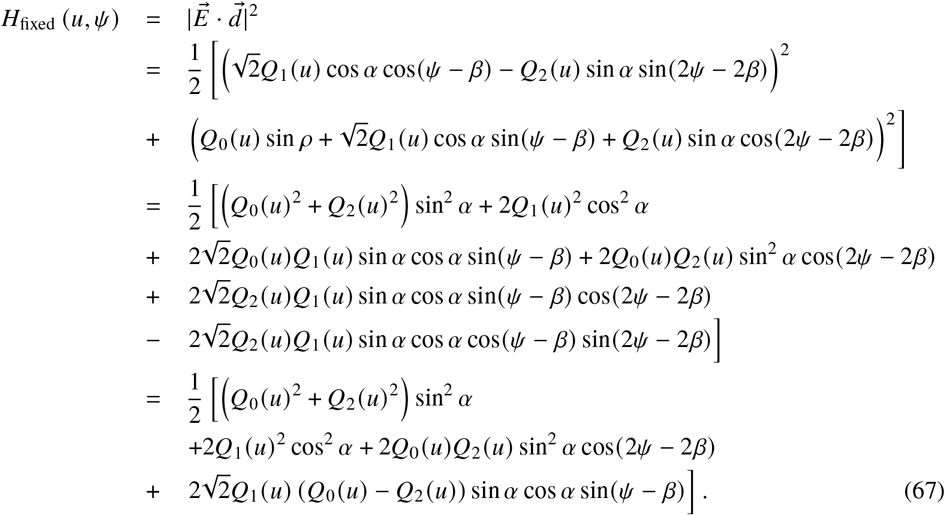

The expressions for the absorption PSF for freely rotating and fixed absorption dipoles are very similar to the expressions for the emission PSF for freely rotating and fixed emission dipoles derived by us earlier [18].

## Appendix B CRLB for Gaussian doughnut PSF model

The Poisson rates for the case where the fluorophore is at the center doughnut position (*x*_0_ = *y*_0_ = 0) are:

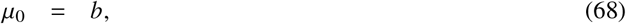

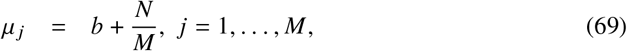

and the *V*_*k*_ are:

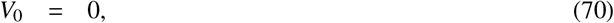

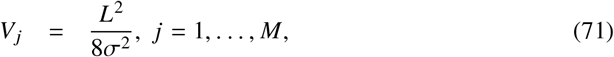

giving:

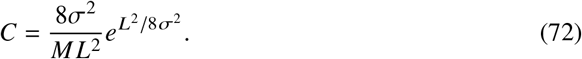

The derivatives of the Poisson rates follow as

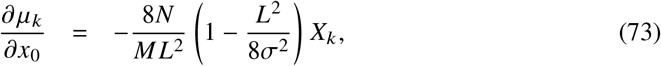

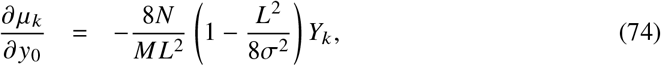

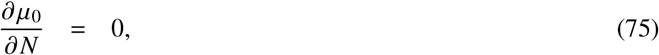

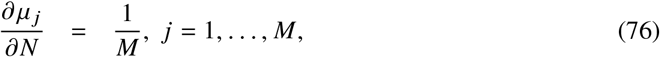

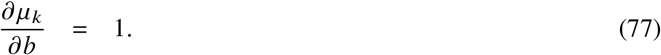

This results in a Fisher matrix:

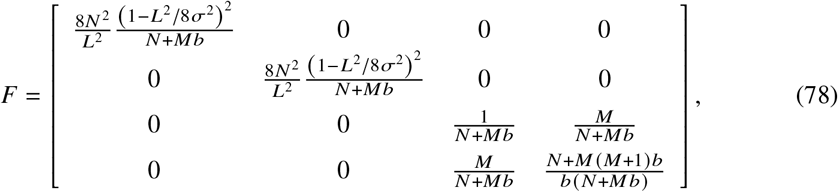

which can be inverted to:

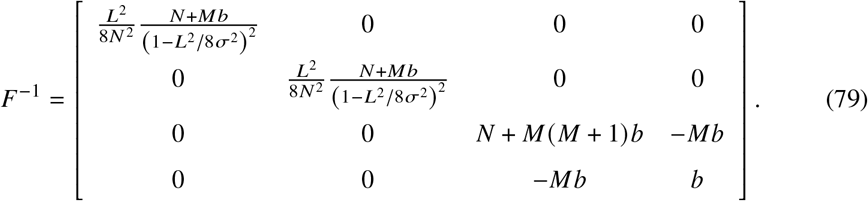

The CRLB follows from the diagonal elements of the inverse Fisher matrix.

## Funding

European Research Council (101055013); Nederlandse Organisatie voor Wetenschappelijk Onderzoek (17046).

## Acknowledgments Disclosures

The authors declare no conflicts of interest.

## Data availability

Matlab code for generating the vectorial doughnut PSF and the MLE fitting is available at https://gitlab.tudelft.nl/imphys/ci/minflux-dipole under Apache 2.0 license.

## Notes

### Competing Interest Statement

The authors have declared no competing interest.

### Summary of Updates

Correct error caption fig. 1; clarifications section 4; correct error fig. 5; expanded discussion fig. 8; added conclusion; new fig. 11; expanded discussion appendix

